# Deactivation and Collective Phasic Muscular Tuning for Pointing Direction: Insights from Machine Learning

**DOI:** 10.1101/2023.03.15.532800

**Authors:** Florian Chambellant, Jeremie Gaveau, Charalambos Papaxanthis, Elizabeth Thomas

## Abstract

Arm movements in our daily lives have to be adjusted for several factors in response to the demands of the environment, for example, speed, direction or distance. Previously, we had shown that arm movement kinematics is optimally tuned to take advantage of gravity effects and minimize muscle effort in various pointing directions and gravity contexts (***Gaveau et al., 2016***). Here we build upon these results and focus on muscular adjustments. We used Machine Learning to analyze the ensemble activities of multiple muscles recorded during pointing in various directions. The advantage of such a technique would be the observation of patterns in collective muscular activity that may not be noticed using univariate statistics. By providing an index of multimuscle activity, the Machine Learning analysis brought to light several features of tuning for pointing direction. In attempting to trace tuning curves, all comparisons were done with respects to pointing in the horizontal, gravity free plane. We demonstrated that tuning for direction does not take place in a uniform fashion but in a modular manner in which some muscle groups play a primary role. The antigravity muscles were more finely tuned to pointing direction than the gravity muscles. Of note, was their tuning during the first half of downward pointing. As the antigravity muscles were deactivated during this phase, it supported the idea that deactivation is not an on-off function but is tuned to pointing direction. Further support for the tuning of the portions of the phasic EMG containing only negative activity was provided by progressively improving classification accuracies with increasing angular distance from the horizontal. Overall, these results show that the motor system tunes muscle commands to exploit gravity effects and reduce muscular effort. It quantitatively demonstrates that phasic EMG negativity is an essential feature of muscle control.

## Introduction

How the brain plans and executes movements is a challenging question in the field of motor neuroscience. The well-known ‘problem of redundancy’ (***Bernshtein, 1967***; ***Latash, 2012***), implies that a large number of joints and muscles must be coordinated to produce adaptive movements that advantageously interact with our environment (***Franklin and Wolpert, 2011***; ***Farshchian et al., 2018***). A fundamental aspect of this environment is the presence of gravity acceleration. Until now, the neural control of movement has been mostly investigated using tasks where the mechanical effects of gravity were canceled (for example, moving in a constrained horizontal plane). However, living organisms must continuously produce adaptable movements that interact with gravity. In attempting to gain knowledge that can apply to real life situations, it is paramount to understand gravity-related movement control. Previously, we demonstrated that the brain plans arm movements whose kinematics are optimally tuned to take advantage of gravity effects and minimize muscle effort (***Gaveau et al., 2016***). Across varied directions and gravity levels, ratios of acceleration versus deceleration well supported our optimal integration of gravity hypothesis and falsified the pre-existing hypothesis according to which the brain would use a tonic torque to compensate for gravity effects (***Hollerbach and Flash, 1982***; ***Atkeson and Hollerbach, 1985***; ***Flanders and Herrmann, 1992***). However, as noticed by reviewers, an important caveat in our investigation and claims was the lack of support from muscle activation patterns (see reviewers comments in the Decision letter for ***Gaveau et al. (2016***)). Since many different muscle contractions can give rise to the same kinematics (***Burdet et al., 2001***; ***Hagen and Valero-Cuevas, 2017***), whether muscle effort truly incorporates the benefits to be obtained from gravity effects remained to be seen.

A qualitative demonstration of a muscular integration of gravity effects for effort minimization was later provided in our study on monkeys and humans (***Gaveau et al., 2021***). Scrutinizing the phasic activations of antigravity muscles i.e. the remaining activity after subtracting the part that compensates for gravity torque (***Flanders and Herrmann, 1992***; ***Flanders et al., 1996***) – we observed that antigravity muscles systematically exhibited negative phases. These negative portions on phasic muscle EMGs indicated that there was less activity in the muscle pattern than what is required to compensate for gravity torque. Importantly, these negative phases happen when gravity is able to help produce the arm motion i.e. during the acceleration of downward movements and during the deceleration of upward movements (***Gaveau et al., 2021***). Analyzing the negativity of phasic muscle activations has since helped to understand the effect of laterality on sensorimotor control (***Poirier et al., 2022***) as well as its modification with increasing age (***Poirier et al., 2023***).

The above studies supporting the optimal integration of gravity hypothesis at the muscular level remain rather coarse-grained. They investigated a limited number of muscles during purely vertical and horizontal movements (***Gaveau et al., 2021***; ***Poirier et al., 2022, 2023***). A better understanding of the integration of gravity in motor control requires a finer-grained investigation where more directions and muscles are examined. In a previous study, we had reported on the kinematics of pointing movements performed in seventeen different directions (***Gaveau et al., 2016***). In this same investigation, we had also recorded the activation patterns of nine muscles for eleven participants but did not exploit the muscle activation results. The present study takes advantage of this EMG dataset to provide a fine-grained analysis of phasic EMGs and examine how they may incorporate the beneficial contribution of gravity for muscle effort minimization. An important caveat in the existing literature is that no study has specifically quantified the importance of the phasic EMG negativity in the overall muscle activation pattern. Whether this negativity represents an important part of the neural signals producing body limb movements is still unknown. In addition, recent reports in the domain of muscle synergies have emphasized the need to develop tools that allow investigating such negative signals (***Scano et al., 2022***; ***Brambilla and Scano, 2022***; ***Scano et al., 2023***).

As mentioned in the previous paragraph, the experiments for the study from ***Gaveau et al. (2016***) were accompanied by the simultaneous recording of muscular activity. As we had recorded EMG activities from 9 muscles of 11 participants performing 12 pointings trials in 17 directions, we ended up with 20196 EMG traces (recorded at 2000Hz). As in the case of many motor control experiments, this constituted a big data base. Consequently, we employed a Machine Learning (ML) technique to draw benefits from a multimuscle approach. By the latter, we refer to situations in which patterns or clearer effects are seen with muscle groups rather than single muscles. The specific approach used was one of binary classification in which EMG time series were automatically classified as coming from a horizontal movement or from another angle. We have previously used binary classification successfully to provide insight into Whole Body Pointing (***Tolambiya et al., 2011, 2012***) and gait (***Nair et al., 2010***; ***Laroche et al., 2014***). Classification accuracy provides an indicator of data separation. Poor separation in the EMGs would lead to poor classification accuracy while EMGs that are very different would lead to higher accuracy. The main Machine Learning classification technique used in the present study was Linear Discriminant Analysis. This algorithm utilizes information on the mean and variance of each category to build a representation of the group (***Grimm and Yarnold, 2000***; ***Johnson and Wichern, 2007***; ***Izenman, 2008***). One of the reasons for choosing LDA is the greater ease for finding the factors which are critical in classification success. This is in keeping with the spirit of explainable artificial intelligence (XAI) (***Murdoch et al., 2019***; ***Arrieta et al., 2020***). Algorithms like Support Vector Machines are more powerful classifiers but harder to probe for the combination of features which are the most critical to classification accuracy (***Thomas et al., 2023***). Nevertheless, as our previous studies have shown that they can be successfully applied to the analysis of EMG data (***Tolambiya et al., 2011, 2012***), we used the technique as a supplementary method to reinforce the conclusions made using LDA.

## Method

### Subject, task and recording

The data used in the current investigation comes from a previous study by ***Gaveau et al. (2016***). To briefly summarize the task: 11 adults were asked to execute a pointing task. Participants started at an initial position (shoulder elevation 90°, shoulder abduction 0°, arm completely stretched) then proceeded to rapidly point at specific directions (every 15°) rotating only the shoulder (see Figure 1a). The EMGs from nine muscles were recorded during the pointing tasks (see Figure 1b). The EMGs were then processed to extract the phasic part of the movement (***Flanders and Herrmann, 1992***; ***Flanders et al., 1996***; ***Gaveau et al., 2016***; ***Poirier et al., 2022***). To do so, EMG signals were integrated from 1 to 0.5s before movement onset and from 0.5 to 1s after movement stop and averaged. The linear interpolation between those two values was used as an estimate of the tonic component, which was removed from the integrated EMG to keep only the phasic component. Then, all the trials were normalized in duration to 1000 points using linear interpolation.

**Figure 1.**
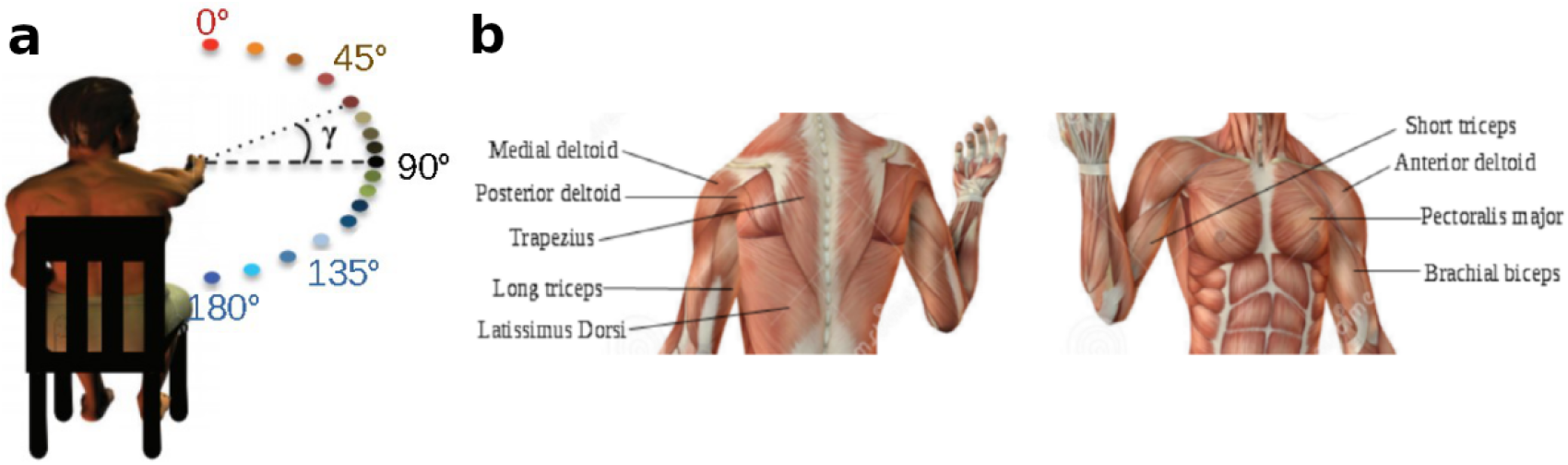
a) Illustration of the task (adapted from ©2016, Gaveau et al.). b) Shoulder muscles from which the EMGs are recorded.

The next step consisted of normalizing EMG amplitude to prepare the input signals for the LDA algorithm. The EMG signals were normalized using a Z-score transformation. It was done separately for each individual. This reduced the variability which would be caused by inter individual differences in EMG amplitudes. It was also done separately for each muscle, hence ensuring that muscles with small EMG amplitudes were not eliminated from playing a role in classification. Finally, a normalization was done for each angle. This normalization removed information on differences in EMG amplitudes for pointing direction. It was an important step as the differences in amplitude could reflect differences in tonic activity for pointing direction rather than provide information on adjustments in phasic activation.

### Linear Discriminant Analysis

Linear discriminant algorithm (LDA) is a classification algorithm which tries to find the best linear hyperplane (hence the name) in the feature space to separate different classes of objects (***Grimm and Yarnold, 2000***; ***Johnson and Wichern, 2007***; ***Izenman, 2008***). This then permits the classification of new observations depending on which side of the hyperplane they are on. Traditionally, LDA was reduced to an eigenvector problem, with the objective of finding a subspace in the original data space that maximize the separability between classes along the first axis of this subspace. However, computational power allows for a more direct approach to classify new data.

To achieve this, the algorithm is used to estimate the probability distributions of the different classes, following certain assumptions about these distributions — that both classes follow Gaussian distributions with the same variance, but different means. Knowing this information, it is then possible to infer the probability of a new observation to belong to a certain class thanks to Bayes’ theorem. This probability is weighted by a cost function to bias the results for particular applications if needed. In practice, the class *ŷ* of a new observation is determined by the following equation:

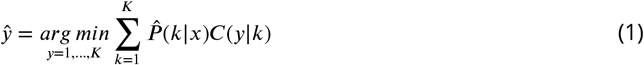

with *y* and *k* being class numbers among the *K* different classes of object.

Thus Equation (1) tries to determine, for an observation *x*, the class *k* that maximizes 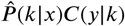 as it is the probability of the observation *x* to belong to class *k*. It is determined thanks to the Bayes’ theorem :

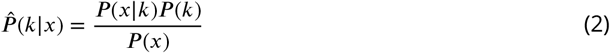

*P* (*k*) is the prior probability and is determined empirically using the labels distribution from the training set. What the algorithm learn from the data are actually the likelihood distributions *P* (*x*|*k*). In the case of LDA, those distributions are assumed to be multivariate Gaussian distributions, all with the same variance matrices. Thus, the main part of the computation is to find the variance matrix and class means. As the distributions are Gaussian, *P* (*x*|*k*) is entirely determined by the Mahalanobis distance of *x* for class *k*. Finally, *P* (*x*) is the normalization factor calculated pooling all the estimated *P* (*x*|*k*) distributions :

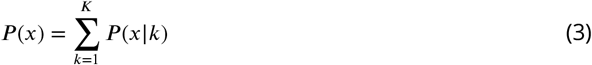

In Equation (1), *C*(*y*|*k*) represent the cost function, *i*.*e*. how much is penalized the misclassification of an observation. This function can be tweaked for particular usage (*e*.*g*. to increase sensitivity of the algorithm at the expanse of specificity). Here, we do not apply a bias in favor of one class, which then translates to the following cost function :

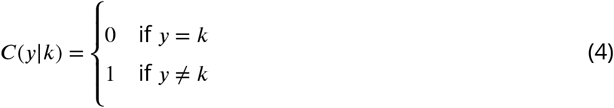

meaning that all classes are equally penalized for misclassification, hence, the hyperplane separating both classes will maximize the separability between the classes and corresponds to the hyperplane that would be obtain through the traditional approach of LDA.

Having all the elements needed, *ŷ* can then be determined for any new observation using Equation (1).

Additionally, as we did in ***Thomas et al. (2023***), we computed the LDA_distance_ as further indicator of the separation of the data. We defined this value as the Euclidean distance between the means of the Gaussian distributions representing the classes

### Support Vector Machine

Like LDA, Support Vector Machines (SVM) are used to find the hyperplane that maximize the separability between two classes (***Cristianini and Shawe-Taylor, 2000***; ***Hastie et al., 2009***). In our study, we used a linear kernel. However, contrary to LDA which takes into account all the sample of a class, SVM is only interested in the samples from a class that are the closest from the samples of the other class (the so-called support vectors). A margin demarcates each class and the objective of SVM is to maximize the separation between these margins. The optimal separation hyperplane is the one in the middle of the largest margin.

This optimal hyperplane can be defined by its orthogonal vector *β* and a bias vector *b* so that :

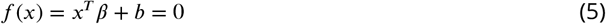

with *x* being an observation. The objective at this point is to minimize ||*β*|| . However, this approach only works if the two classes are perfectly separable, which may not be the case here, hence we introduce so-called ‘slack variables’ which will penalize the margin size for each support vector on the wrong side of the separation hyperplane. Thus, the objective becomes to minimize :

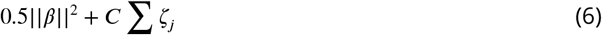

with respect to *β, b* and *ζ*_*j*_ subject to *y*_*i*_*f* (*x*_*j*_) ≥ 1 − *ζ*_*j*_ and *ζ*_*j*_ ≥ 0 for all *j* = 1, …, *n. y*_*j*_ is the label of the observation *j* (either −1 or 1) and *C* is a positive scalar called ‘box constraint’ that regularizes the penalty assigned to errors. To optimize the objective function presented in Equation (6), we use the Lagrange multipliers method. This method introduce a series of coefficients *α*_1_, …, *α*_*n*_ which allow the transformation of the objective into maximizing :

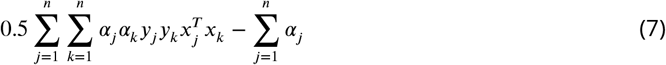

with respect to *α*_1_, …, *α*_*n*_ subject to :

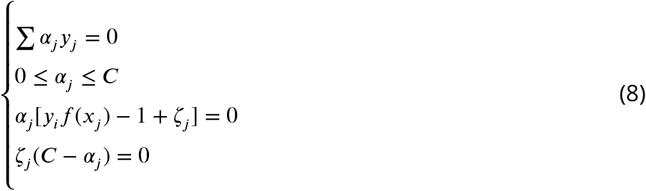

for all *j* = 1, …, *n*. Finally, a new observation *z* is assigned to a class following the equation :

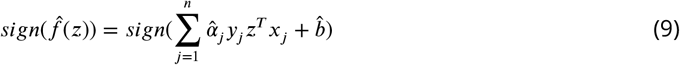

where 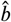 is the estimate of the bias and 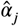 is the *j*^*th*^ estimate of the vector 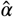. Once classification was done, we used 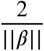 as an estimate of the size of the margin separating the data.

### Data Organization for Machine Learning

All analyses were performed offline using custom Matlab scripts (***MATLAB, 2022***). The input data was constructed by concatenating the processed EMG signals of the nine muscles. This gave us one vector of 9000 points (1000 normalized time points multiplied by 9 muscles) per trial. Thus, the algorithms were trained on the entire EMG waveforms of all the muscles simultaneously. Further analyses involved focusing on certain parts of the signal. Portions of this vector were isolated as necessary when analyses only involved some muscle subsets (e.g. only the gravity muscles).

The training of the algorithms was done using binary classification i.e. the algorithm was trained to discriminate between EMGs of two directions. One of the movement directions was always the horizontal plane (90°, gravity neutral, taken as reference) while the other class was any of the other angles.

To ensure the generalization of the results, a stratified five-fold cross-validation method was used. The data set was separated into training and testing sets using the known labels of the samples to ensure that both directions were equally represented in each training and testing set. The cross-validation approach allowed a better estimation of the efficacy of the algorithm by testing it on multiple data sets. Thus, we were able compute an average accuracy of the algorithm on several testing sets.

From each trained LDA algorithm, we extracted the means of the two *P* (*x*|*k*) distributions as well as the testing accuracy. Then, we computed the LDA_distance_ measuring the separation between the mean vectors of each class. For SVM algorithms, we extracted the vector orthogonal to the separation hyperplane and computed the width of the margin.

### Statistical analysis

The significance of results was obtained using non-parametric tests, namely the Wilcoxon rank sum test when comparing two conditions and the Friedman test to compare effects across multiple conditions. Results were considered to be significant for *p* < 0.05.

## Results

The processed phasic EMGs from the recorded nine muscles for every pointing angle can be seen in Figure 2. We will start the result section with a presentation of a grid displaying the classification accuracies of individual muscles versus the accuracy obtained from all the muscles. The main purpose of this would be to illustrate the higher accuracies obtained from using the combined muscle information. We will then continue examining the classification obtained using all the muscles simultaneously in order to trace the nature of collective muscular tuning for pointing direction. The nature of this tuning will be further analyzed using the LDA_distance_. From using input vectors which contain the information from all the muscles, we will then turn to analyzing specific groups of muscles. Once again, classification accuracies will be used as an indicator of adaptation for pointing direction, hence allowing us to trace tuning curves for particular muscle groups. Finally, as one of the major points of this paper is that the negative portions of the phasic EMGs are a result of gravity effort optimization, we will use classification to see if this portion of the EMG is tuned to pointing direction. Results obtained using LDA will also be reinforced using the SVM as an alternate Machine Learning algorithm.

**Figure 2.**
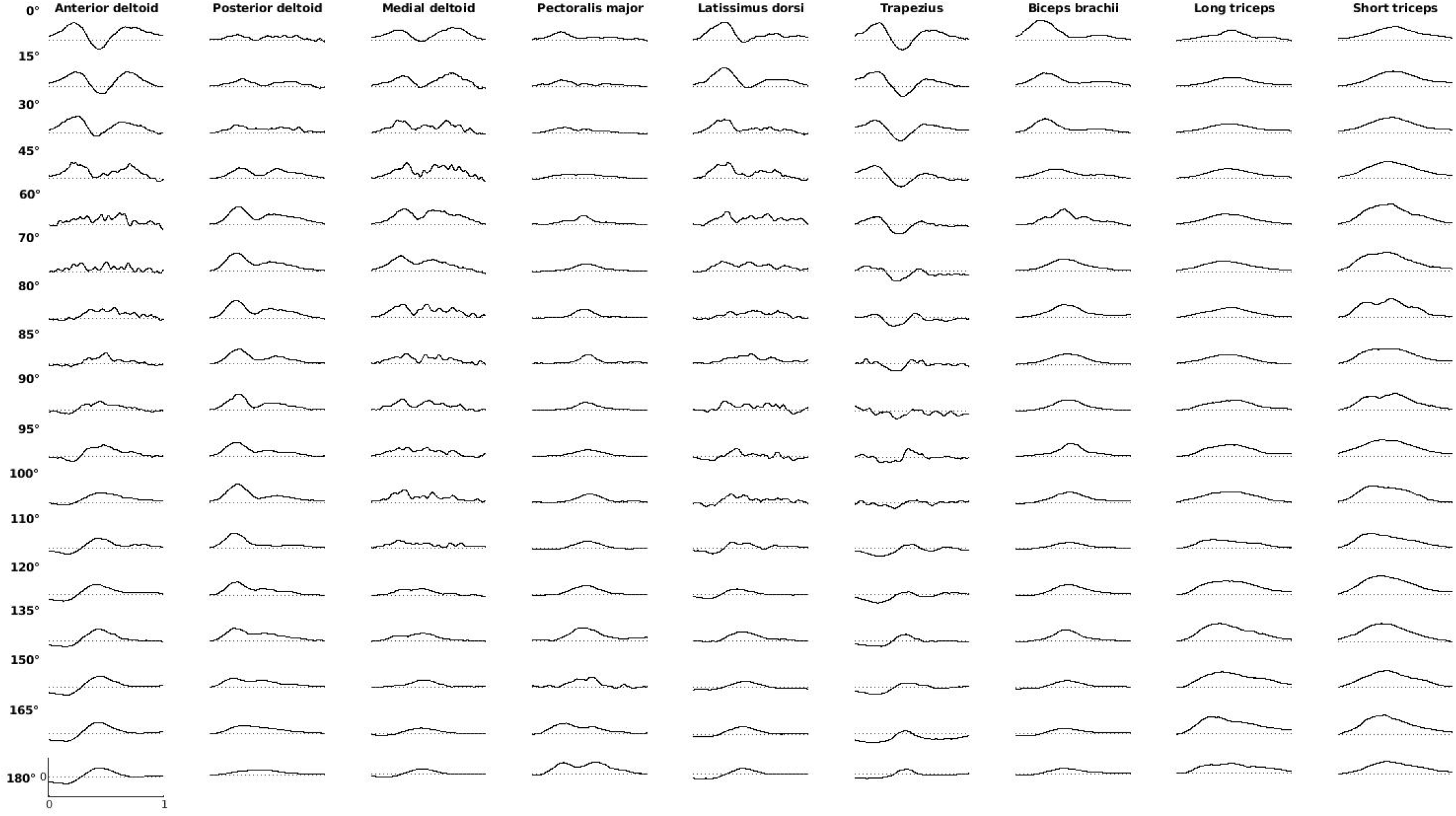
Example of EMG recordings (averaged among all trials from one participant). Data was normalized by the highest recorded phasic activity of the participant.

### Advantage of a multimuscle approach

The aim of this section is to demonstrate some of the benefits of a multimuscle approach. In Figure 3 we display a grid with the classification accuracies of individual muscles for each pointing angle as well as the accuracy using all the muscles simultaneously (last line of grid). For almost all directions, the classification accuracy using all the muscles is higher than those obtained with individual muscles. This demonstrates that information concerning differences in muscular activity for pointing in different directions may be found at the level of the muscle population even when it may not be significantly present at the level of individual muscles.

**Figure 3.**
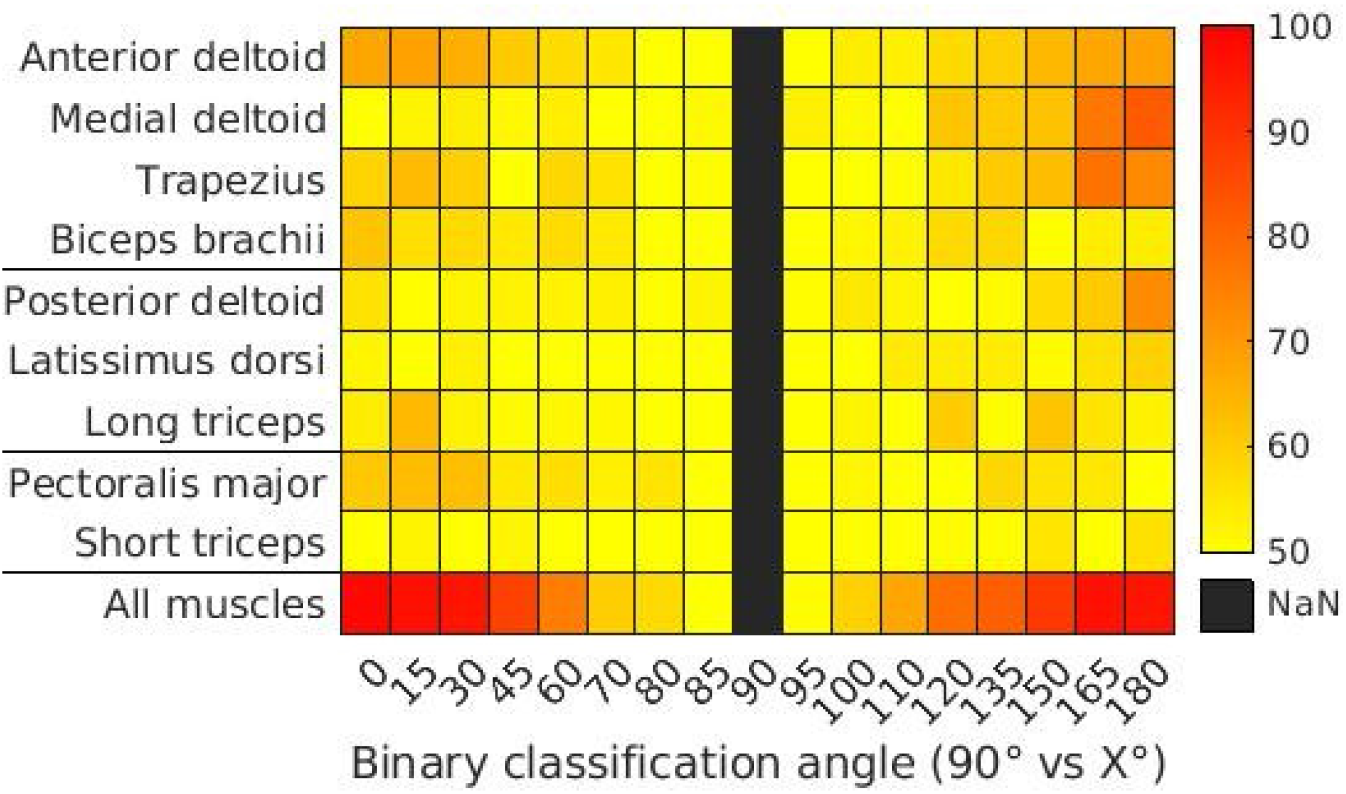
Average binary accuracy of individual and all muscles for different angles compared to 90°, using LDA for classification. The muscles are ordered by their type (anti-gravity, gravity and neutral) and lines separate the different types of muscles.

Other features of note in this figure are that some muscles like the anterior deltoid, have relatively high accuracies when pointing at 0° as well as at 180°, hence indicating an important role in adaptation for both directions. This is not the case for a muscle like the medial deltoid which displays high accuracies for pointing at 180°, but not for 0°. The short triceps has low accuracies in discriminating for any angle, hence showing that it does not have a prominent role in adapting for direction.

### Binary classification with all muscles for pointing direction

The aim of this section is to follow the nature of multimuscle adaptation for pointing in each of the seventeen directions described in the Methods section. Using all the muscles, LDA was used to detect if pointing had taken place horizontally or at another pointing direction. Figure 4 displays the accuracies from this classification of 90° (horizontal pointing) versus another angle displayed on the x-axis. Poor classification accuracies reflect small separation between data groups while a good accuracy is indicative of clear separation. The EMG tuning curves using this technique display adjustments which are mostly linear until approximately 60° from horizontal direction. Following this, classification accuracies took a non-linear trend. Similar trends were observed when performing the same classification with the SVM (Supplementary Figure S1).

**Figure 4.**
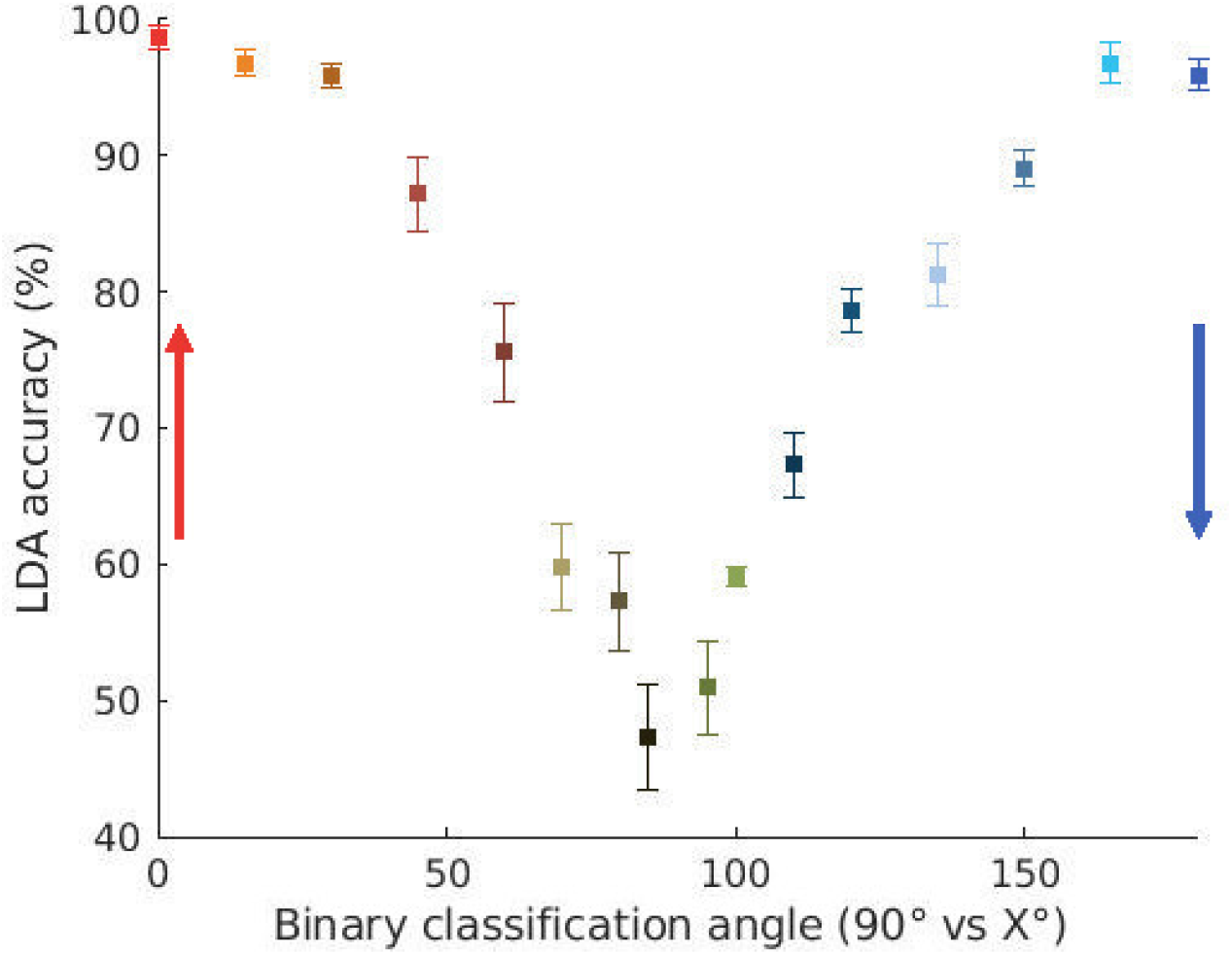
Accuracy of the binary classification between 90° and the other angles, using the EMGs from all muscles for LDA classification. Error bars indicate standard error. Arrows indicate pointing directions considering as origin the horizontal axis (90°). The colors indicate pointing directions as in Figure 1a.

The non-linearity in binary classification may not be due to actual non-linear muscular modification but due to saturation in classification capacities (see Supplementary Figure S9). Verification as to whether this was the case, was done by further examining the model used by LDA to perform the classification.

The multivariate mean of each category is an important piece of information used by LDA to create representations of each category. The distance between these means can be exploited to measure muscular tuning for pointing direction. Figure 5 displays how this distance changes as a function of pointing direction. The figure confirms a linear trend in collective muscular tuning for pointing direction until about 60° from horizontal. However, unlike classification accuracy, LDA_distance_ is not subject to the effects of classification saturation. Muscular modification as reflected in the LDA_distance_s continue increasing with a slightly nonlinear trend both upwards to 0° and downwards to 180°.

**Figure 5.**
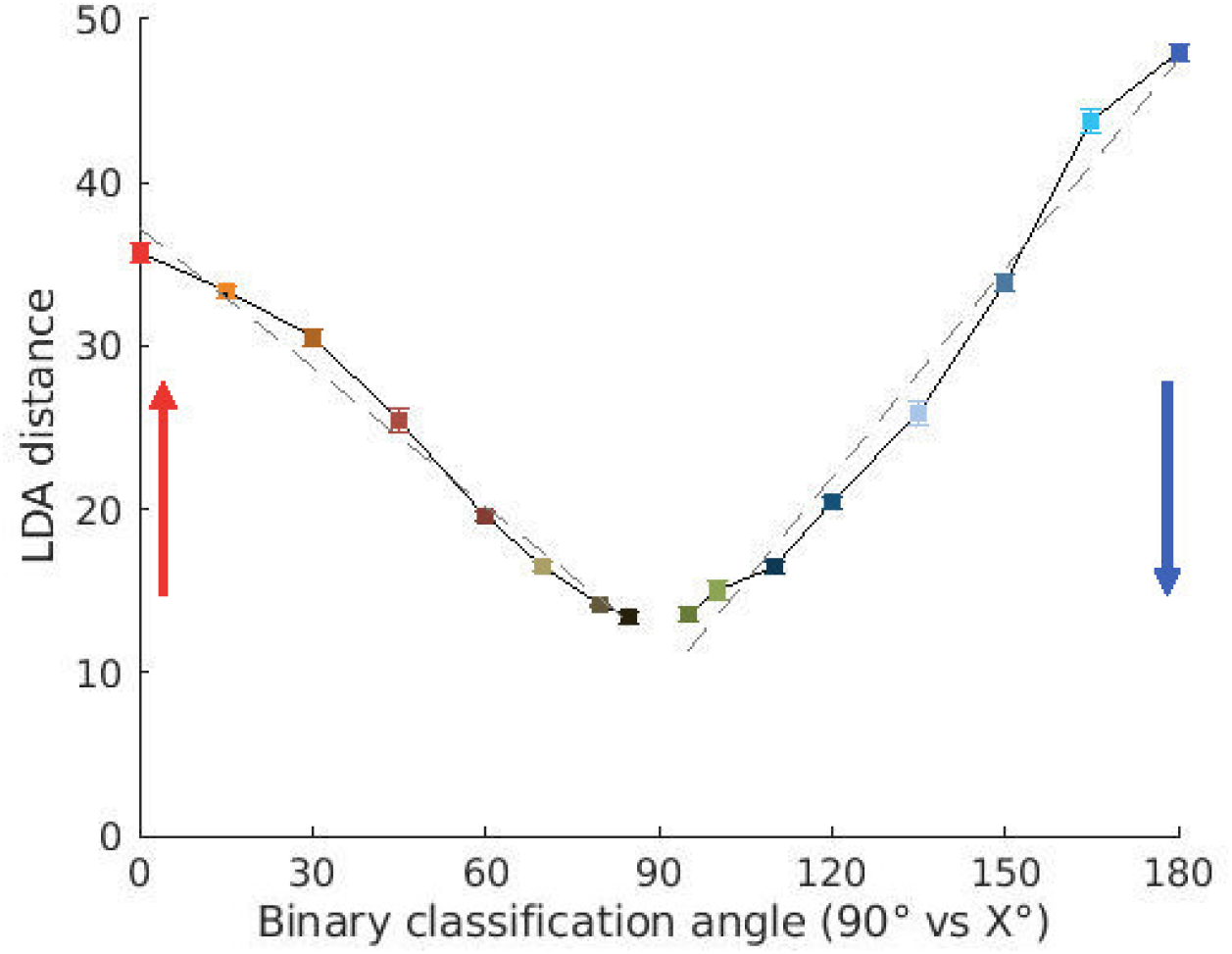
LDA_distance_ computed from the models of each category constructed by the classification algorithm for binary discrimination between 90° and the other angles using all muscles (see Methods). Error bars indicate standard error. Gray dashed lines show the linear regressions performed for comparing upward versus downward tuning. The colors indicate pointing directions as in Figure 1a.

The LDA_distance_ also revealed an intriguing aspect of muscular tuning for pointing direction. It showed that adjustments for downward pointing are bigger than those for upward pointing when compared to pointing in the horizontal direction. The LDA_distance_ for pointing at 0° upward was found to be significantly lower than the distance for pointing at 180° downward (Wilcoxon rank sum *W* = 0, *p* = 0, 008). Differences in the gain for tuning were also revealed by fitting a regression line for both sides of the EMG tuning curves. They revealed a value of 0.2839 (*R*^2^ = 0.9852) for upward pointing and 0.4257 (*R*^2^ = 0.9795) for downward movements indicating once again that counter to intuition, movements were more finely tuned in the downward direction. Further confirmation of this difference in upward and downward adaptation can be seen in the Supplementary Figure S2 where we measure the distance from both groups using another classification algorithm, the Support Vector Machine (SVM, Wilcoxon rank sum *W* = 0, *p* = 0, 008).

### Modular EMG tuning for pointing direction

As the system adapts for tuning in various directions, what is the nature of this tuning? Does the entire system undergo modifications to bring the arm to a new direction i.e. a uniform tuning? Or is adaptation for pointing to the new direction primarily accomplished by a few muscles at particular moments, i.e. modular tuning? To answer this question, we examined the classification capacities of different groups of muscle at different moments. The muscle groups were the gravity and antigravity muscles. A finer grained analysis of modularity was obtained by examining the acceleration and deceleration halves for the aforementioned muscle groups. The gravity muscles in this study are the posterior deltoid, the latissimus dorsi and long triceps. The antigravity muscles are the anterior deltoid, medial deltoid, trapezius and biceps brachii. Figure 6 below demonstrates the classification accuracy of these muscle groupings. Taking the case of the acceleration phase it can be seen that the antigravity muscles achieved a significantly superior performance at predicting pointing direction when compared to the gravity muscles (Friedman test, *χ*^2^(1) = 26.35, *p* < 0.001).

**Figure 6.**
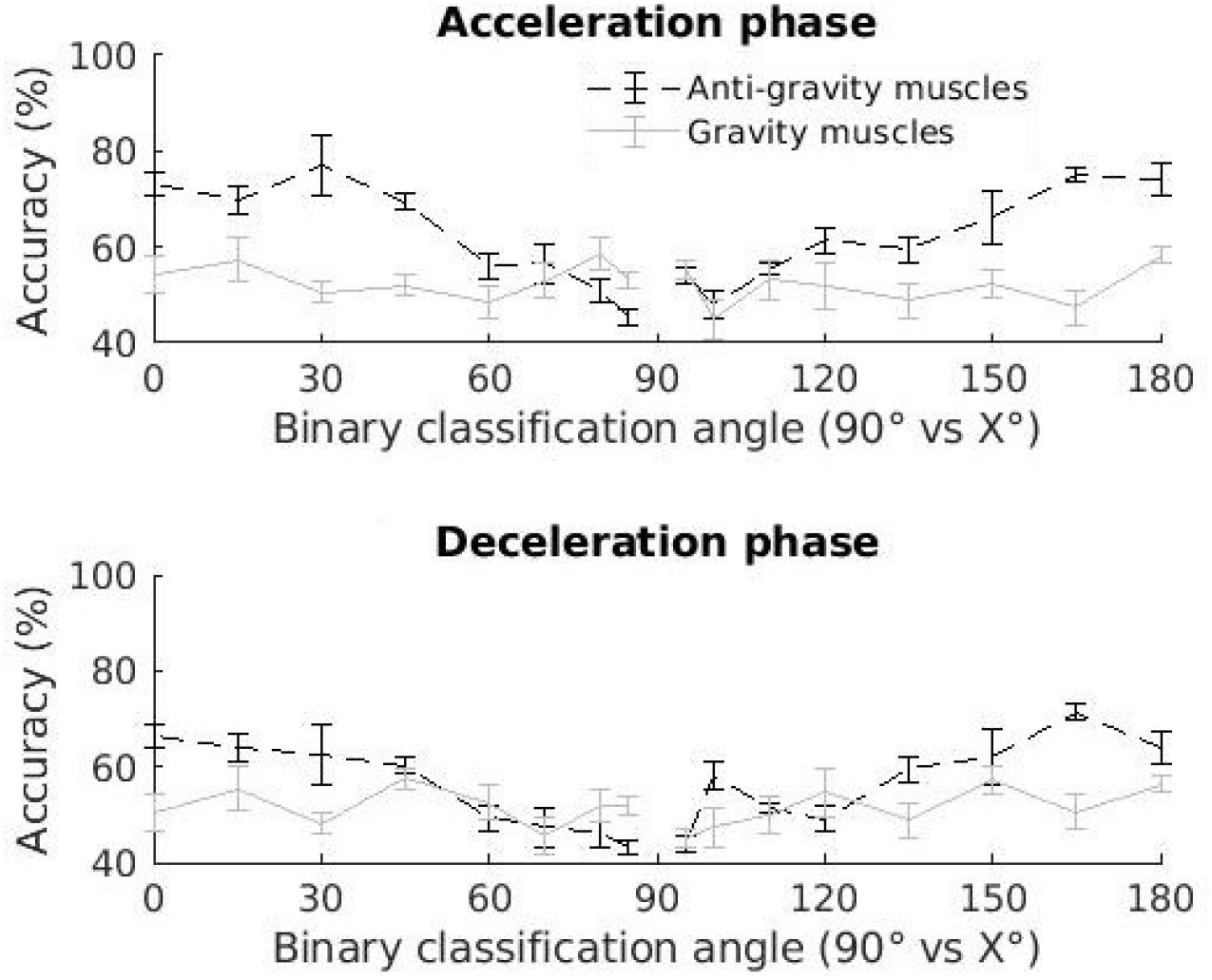
Classification accuracy using only the EMGs from either gravity or antigravity muscles during the acceleration or deceleration phase of the movement, using LDA for classification.

As there were 4 antigravity muscles but only 3 gravity muscles, it was necessary to verify that the higher classification accuracies observed in the antigravity muscles was not due to more information. To do this, we repeated the classification tests after having removed the biceps brachii from the group of antigravity muscles (this muscle showed the poorest tuning among the group of antigravity muscles, see Figure 3). The prediction capacities of the antigravity muscles remain significantly higher than those of the gravity muscles (Friedman test, *χ*^2^(1) = 10.39, *p* = 0.001, see Supplementary Figure S4).

Table 1a shows the values for the slopes of these lines for each group of muscles for upward and downward pointing in Figure 6. They reveal higher slopes for the antigravity muscles at all phases of upward and downward pointing. Of note, are the slopes of the antigravity muscles during the acceleration phase of downward movement and the deceleration phase of upward movement. As these are phases during which the antigravity muscles have lowered activity, it indicates that the deactivation in these muscles is not an all-or-nothing function, but one that is graduated as a function of pointing direction. The lower values of the slopes for the gravity muscles demonstrate relatively smaller adjustments by this group of muscles for pointing angles.

**Table 1.** Slope coefficients of the linear regressions for the classification accuracy of the different muscle groups from Figure 6.

|  | Upward movements | Downward movements |
| --- | --- | --- |
| Antigravity muscles | 0.3306 ($R^2 = 0.80$) | 0.2910 ($R^2 = 0.90$) |
| Gravity muscles | 0.0041 ($R^2 = 0.001$) | 0.0315 ($R^2 = 0.051$) |

|  | Upward movements | Downward movements |
| --- | --- | --- |
| Antigravity muscles | 0.2895 ($R^2 = 0.93$) | 0.233 ($R^2 = 0.67$) |
| Gravity muscles | 0.0183 ($R^2 = 0.02$) | 0.0983 ($R^2 = 0.49$) |

All these results were further confirmed using another classification algorithm, the SVM. These results can be seen in Supplementary Figure S5.

### Tuning with the negative portions of the antigravity muscles

An important tenet of the gravity effort-optimization hypothesis is that the muscles are able to reduce their activity in conditions where gravity is able to replace their role. In the case of EMG phasic activity, this would especially be seen during the negative phases of this activity, primarily at the latter half of upward movement (deceleration) and the first half of downward movement (acceleration). Figure 6 which examines tuning in the group of antigravity muscles for the acceleration and deceleration halves of pointing already hints at the likelihood of tuning in this phase. Nevertheless, the acceleration and deceleration phases of the antigravity muscles still contain portions of the EMG which are positive. In this section we zone in further to examine the tuning capacities of the portions of the phasic EMGs for the antigravity muscles which are exclusively negative. Figure 7 contains the results of these tests. Once again, the increasing classification accuracy of this portion of the phasic EMG as a function of pointing direction, indicates adjustments as a function of pointing direction. Further confirmation of tuning in this negative portion of the EMG was obtained using the SVM classification algorithm (Supplementary Figure S7).

**Figure 7.**
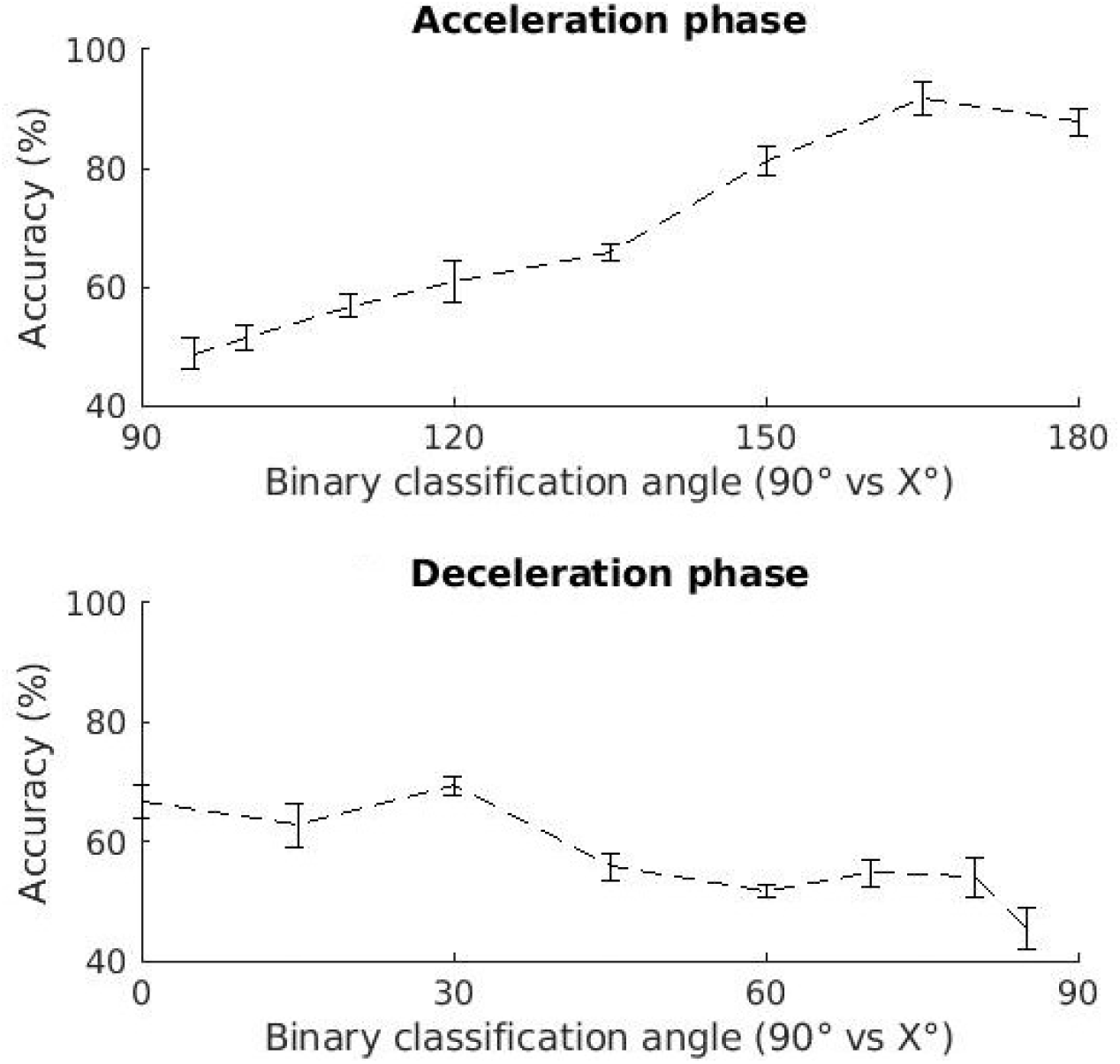
Classification accuracy using only the negative part of the EMGs from antigravity muscles during the acceleration or deceleration phase of the movement, using LDA for classification.

## Discussion

In this study we examined the manner in which phasic activity of shoulder muscles are adjusted for arm pointing movements in different directions. This extends previous results demonstrating that arm kinematics is tuned to pointing direction (***Gaveau et al., 2016***). The researchers had demonstrated that effort optimization leads to clear differences in phasic muscular activity for vertically upward and downward pointing. The authors had also showed qualitatively that this integration can be seen at the muscular level (***Gaveau et al., 2021***). The current study extends on these two previous studies by providing a much finer grained, quantitative picture of phasic muscular tuning for pointing direction. More specifically, through the use of relatively recent quantitative tools, it analyzes the information content in the negative portions of the phasic EMG.

A key finding of the study is that the negative portion of the phasic EMGs contains information concerning pointing direction. Many previous studies had set aside this part of the EMG as unimportant and the inevitable result of the computation required to extract phasic activity (***d’Avella et al., 2008***; ***Russo et al., 2014***). The presence of information in this section of the EMG is especially clear from Figure 7 (and Supplementary Figure S7) where Machine Learning was used to automatically detect if the unlabeled negative portions of the EMGs of antigravity muscles could be used to predict whether a subject had performed horizontal pointing or pointing at some other specified angle. The figure shows that in many cases, accuracy was above chance levels. It also shows that the classification accuracy changed as a function of pointing direction. As explained in the Introduction section, classification accuracy is an indicator of data separability. The increasing classification accuracy with increasing angular separation from horizontal indicates that the deactivation in these negative phases were not on-off functions but scaled in accordance with pointing angle. An on-off deactivation function that does not change with pointing angle would have given similar classification accuracies (perhaps above chance) for all pointing angles. A better idea of the information content in the negative portions of the phasic EMGs can also be obtained by examining the classification accuracies obtained using the positive portions of the phasic EMGs (Supplementary Figure S8). It can be see here that the tuning curves do not attain classification accuracies that are higher than those of the negative portions.

Another original aspect of the study is the use of Machine Learning in order to obtain an ensemble view of muscle activation patterns. Tuning of muscles for different pointing directions was analysed in terms of classification accuracy. The prediction of task constraints using Machine Learning algorithm was done with combined muscular activity as input. This approach is interesting in view of the synergistic manner in which muscles achieve task goals. The term synergistic refers to the fact that movement is the result of the combined activities of several muscles in which the role of any one muscle in the group need not remain constant as others can compensate for this inconsistency and still achieve the original goal. The term Motor Equivalence also refers to this idea (***Lashley, 1933***; ***Morasso, 2022***). An ensemble analysis therefore has the potential to provide insight into group properties which may not be present at the single muscle level or worse yet, at the level of a single EMG parameter such as amplitude or onset delay. A single variable of this sort may not have sufficient power to reach statistical significance. This is clearly demonstrated in Figure 3 where the last line displays the classification accuracy obtained from the combination of all the muscles recorded during the experiments. It is darker than any of the preceding lines hence indicating a higher discrimination power for the muscle population.

The main machine learning technique used for the study was LDA. This technique was primarily chosen because of its simplicity and ease of understanding the features which are critical to the classification (***Thomas et al., 2023***). As a technique which is not as powerful as methods which were developed later (***Statnikov et al., 2008***; ***Nair et al., 2010***; ***Heung et al., 2016***; ***Han et al., 2018***; ***Uddin et al., 2019***), there was also a lower risk of classification saturation (see Supplementary Figure S9). By this we mean high classification accuracies from very small differences, leading to poor information concerning differences in data separability for different pointing angles. We address this problem in Figure 5 where LDA_distance_ was used to probe data separation in situations where classification accuracy had reached a plateau of close to 100%. Our fears on this question turned out to be unfounded when we used the SVM with a linear kernel for the supplementary figures of the current study (Supplementary Figures S1, S2, S5, S6, S7). True to its reputation as a more powerful classifier, the SVM did indeed provide generally higher accuracies but displayed tuning curves with patterns similar to those obtained using the LDA algorithm, hence backing up the results obtained with LDA.

Figure 5 (and Supplementary Figure S2) presents the intriguing result that the EMG separation is higher in the downward direction than upward. One possible explanation of this may be that both upward and horizontal pointing follow the classical triphasic burst pattern (***Hallett et al., 1975***; ***Virji-Babul et al., 1994***). In contrast to this, Gaveau et al had predicted (2016) and demonstrated (2021) a very different pattern of muscular activity for downward pointing. In accordance with the effort optimisation hypothesis they showed that the antigravity muscles phasic activity becomes negative during the first half of downward pointing, hence leading to an activity pattern which is quite different from what is seen for horizontal pointing.

A similar explanation could be used to explain the differences in the tuning curves of Figure 6 (and Supplementary Figure S5). The tuning curves of these figures and Table 1 display a sharper tuning for pointing direction in the antigravity muscles than for the gravity muscles (higher slopes in accuracy as a function of pointing direction). This difference in tuning may be due to the fact that the negative portions of the phasic EMGs are not present in the gravity muscles. The poorer tuning of the gravity muscles may be due to this absence and hence underscores the important role played by the negative portions of the phasic EMGs in adjusting for pointing direction.

Figure 3 displays the accuracies of each muscle for pointing in different directions individually. The accuracy patterns of the anterior deltoid show that it is tuned in both directions. The trapezius shows a similar pattern of classification accuracies, though their smaller values would indicate smaller adjustments than the anterior deltoid. In contrast to the two aforementioned muscles, the phasic activity of the medial and posterior deltoids are more tuned for downward pointing, showing higher accuracies for distinguishing downward than upward pointing angles. The importance of all these muscles in pointing have been highlighted by various studies (***Flanders, 1991***; ***Flanders et al., 1994, 1996***; ***Mira et al., 2021***; ***Tokuda et al., 2016***). In contrast to the previous studies, the use of classification accuracy allows for a greater ease in comparing and contrasting the roles of individual muscles. A direct comparison with the previous studies is not possible, as the pointing protocols were often different, involving for example, mobility around the elbow joint. In contrast to the previously mentioned muscles, the short triceps displays poor classification accuracies at all angles in both directions. Once again, this does not necessarily mean that the muscle is not active, but that it is poorly distinguishable from its activity for horizontal pointing, and hence does not play an important role in tuning for direction. This is not unexpected as this mono-articular muscle is involved in keeping the elbow joint extended but not in rotating the shoulder joint.

No discussion on ensemble methods would be complete without talking about matrix factorization methods. These techniques have been very useful in demonstrating that the panoply of EMG recordings from the arm during pointing under different constraints can be simplified by using a small number of basis functions which can then be adapted for multiple constraints (***d’Avella et al., 2008***; ***Muceli et al., 2010***; ***d’Avella and Lacquaniti, 2013***). The technique which was frequently applied was non negative matrix factorization (NNMF). Therein lies the problem for analyzing a portion of the phasic EMG which this study has found to be important – the negative portion. Since it is a technique with non-negativity constraints, NNMF is not adapted to the study of the negative portions of the phasic EMG. This problem has recently been addressed with the mixed matrix factorization method (MMF) (***Scano et al., 2022, 2023***). Even though, like Machine Learning, these matrix factorization methods are ensemble methods taking into account global properties of big collections of data, they have a very different focus when compared to Machine Learning classification. Their emphasis is on finding commonalities between the collection of EMG trajectories while Machine Learning classification centers on finding differences. The two methods are therefore complementary to each other. It should be noted that in many fields, the two methods are sometimes used sequentially. Matrix factorization is often used in the feature extraction step to first reduce spatial dimensionality before then going on to apply Machine Learning (***Duda, 2000***). This step was not taken in our study as it would have complicated the process of trying to understand the features which were critical to classification.

In conclusion, we will say that Machine Learning classification shows that the antigravity muscles are better tuned to pointing direction than the gravity muscles. Focusing on the deactivation portions of these phasic EMGs, our study demonstrates that they are modified for pointing direction. Previous studies by ***Gaveau et al. (2021***) supported the hypothesis that this negativity would result from integrating gravity for optimized motor control. They had not however, studied the nature of this muscular adjustment in fine detail. Using the EMG patterns from nine muscles and pointing in 17 directions, the current study shows that the deactivation of the antigravity muscles is not an on-off function, but is adjusted as a function of pointing direction.

## Supplementary Material

**Figure S1.**
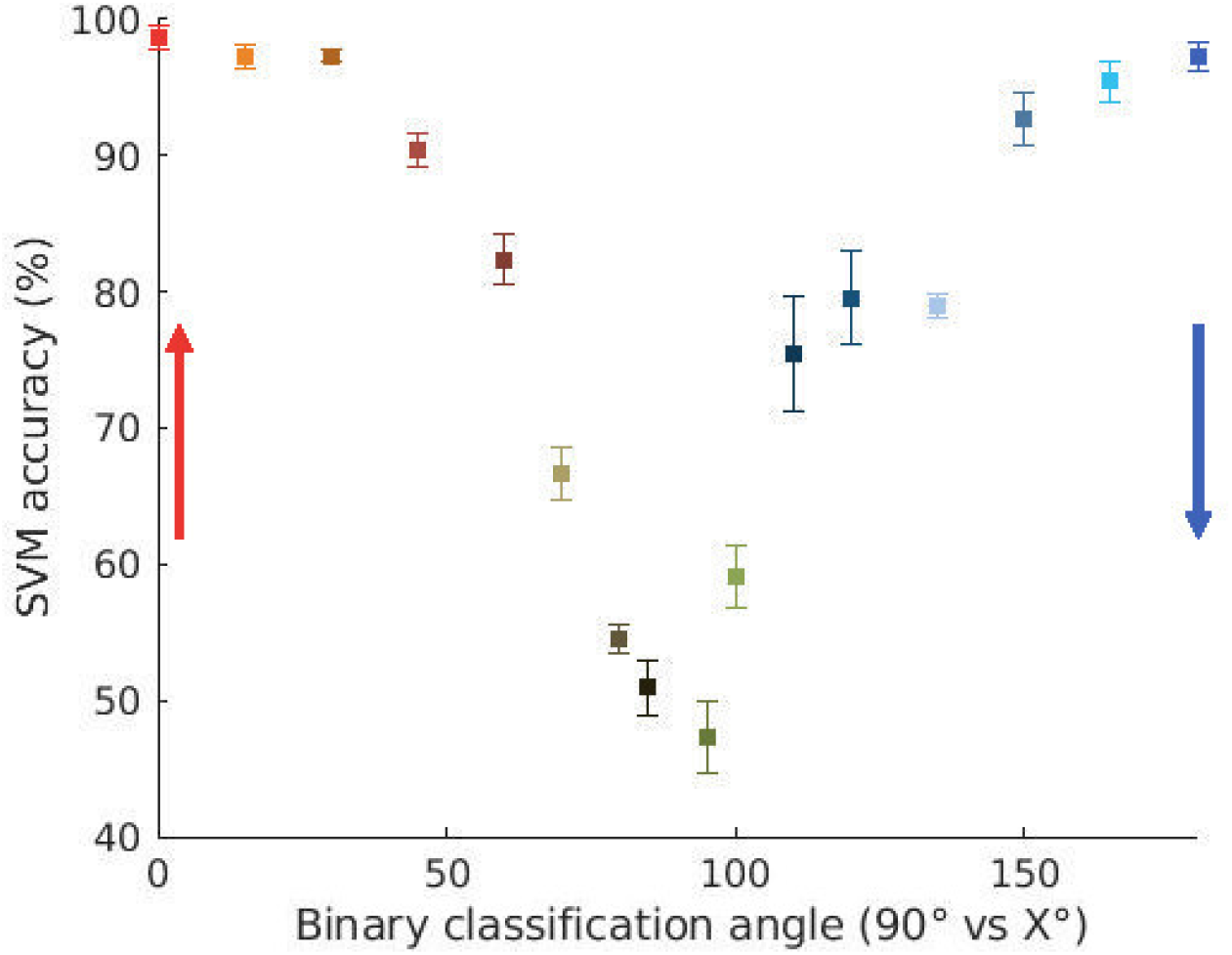
Accuracy of the binary classification when using EMGs from all the muscles. Classification was done for pointing between between 90° and the other angles, using SVM for classification. Error bars indicate standard error. The colors indicate pointing directions as in Figure 1a.

**Figure S2.**
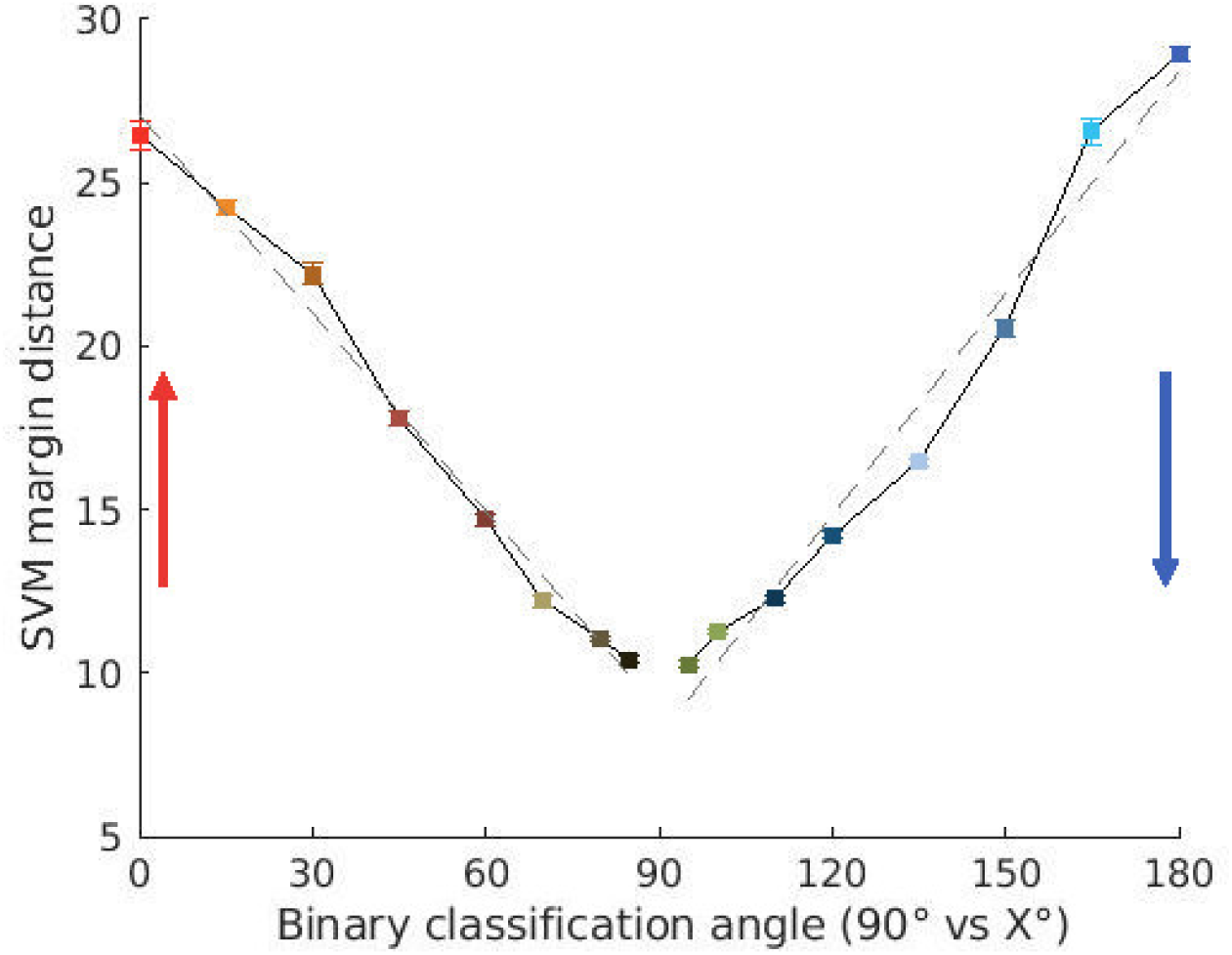
Size of the estimated SVM margin using EMGs from all muscles. Error bars indicate standard error. The colors indicate pointing directions as in Figure 1a.

**Figure S3.**
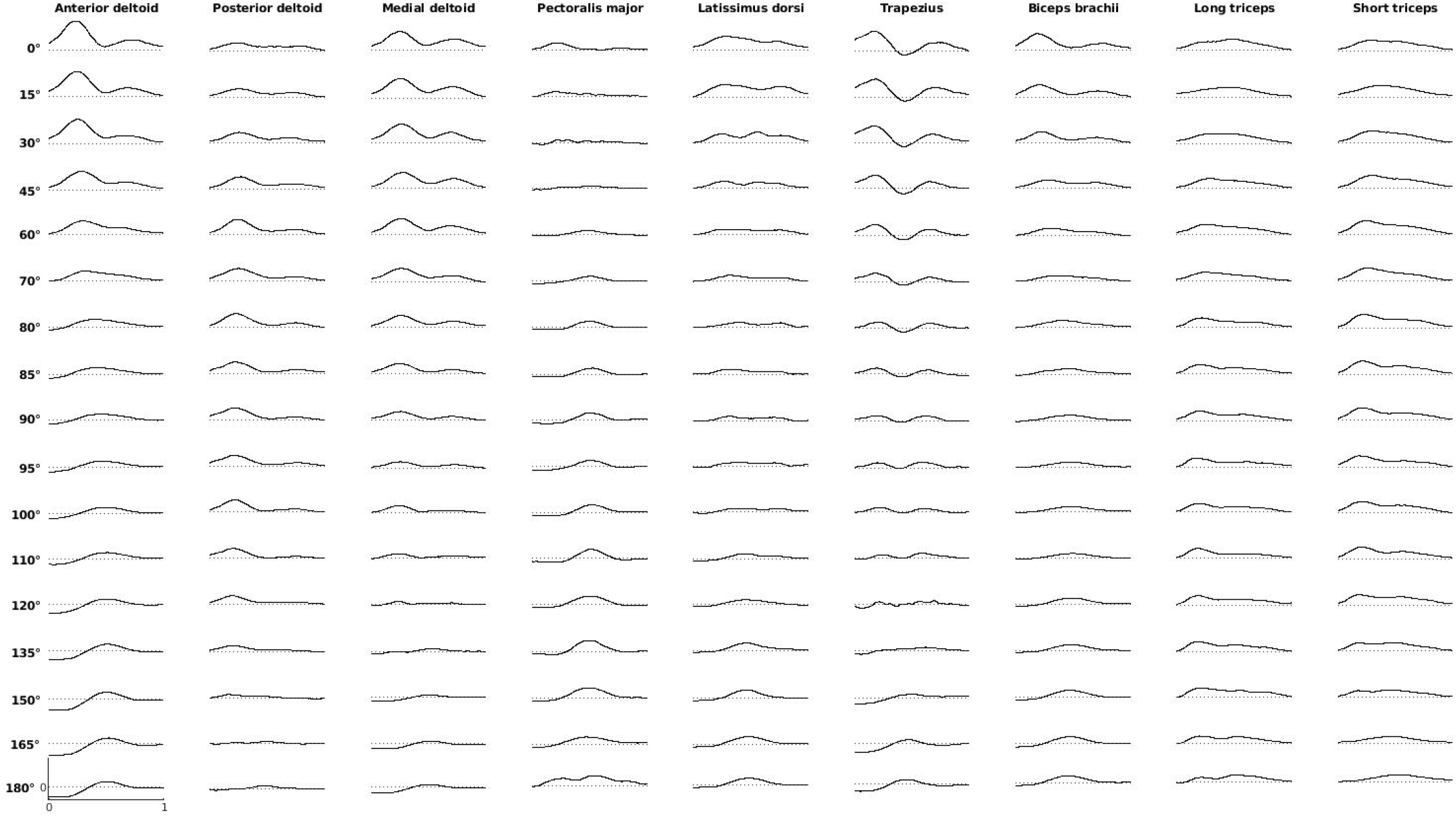
Average of the EMG recordings from all participants. Data was normalized by the highest recorded phasic activity of each participant.

**Figure S4.**
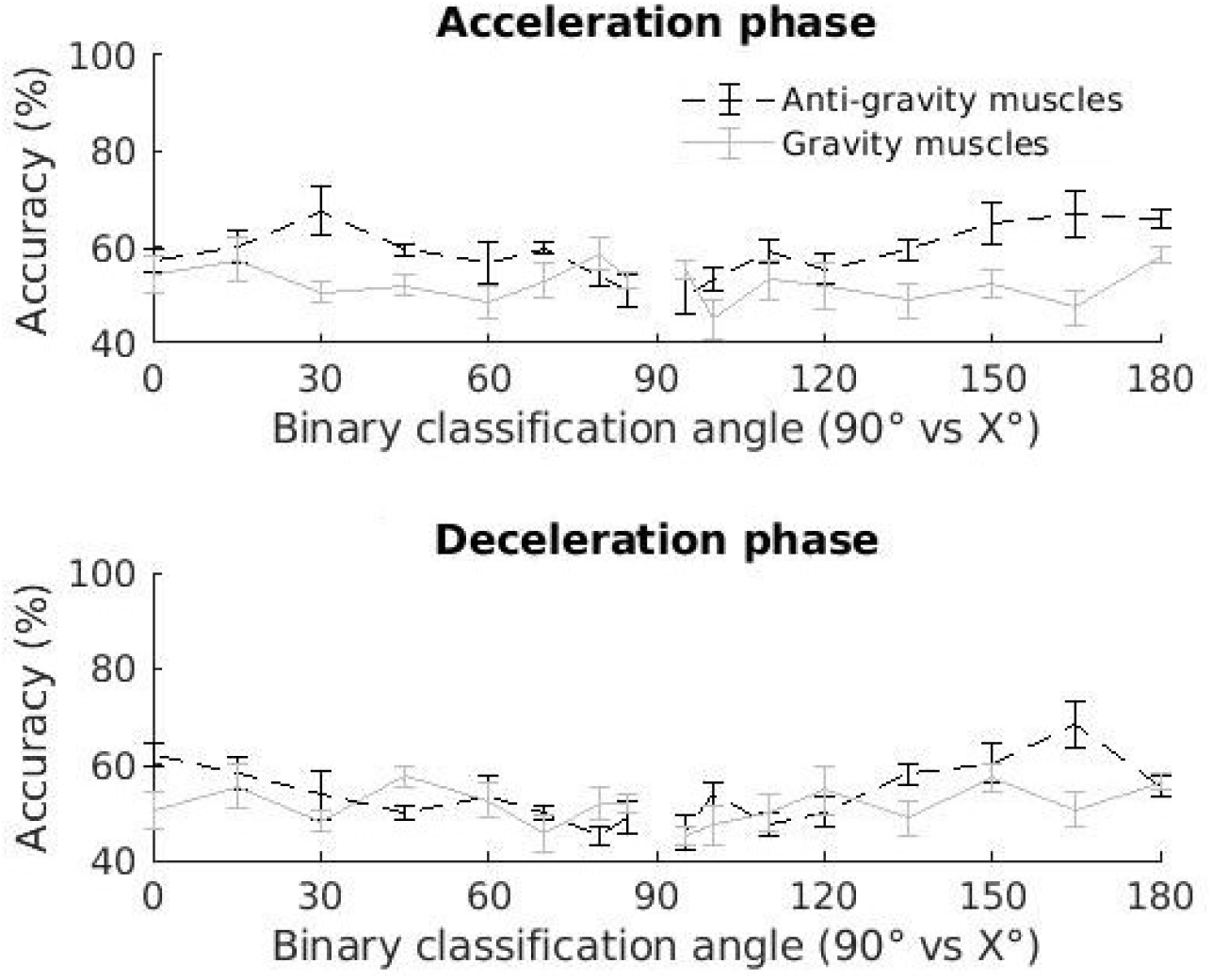
Classification accuracy using only the EMGs from either gravity or antigravity muscles. Contrary to Figure 6, the biceps brachii is not included in the antigravity muscles here, using LDA for classification.

**Figure S5.**
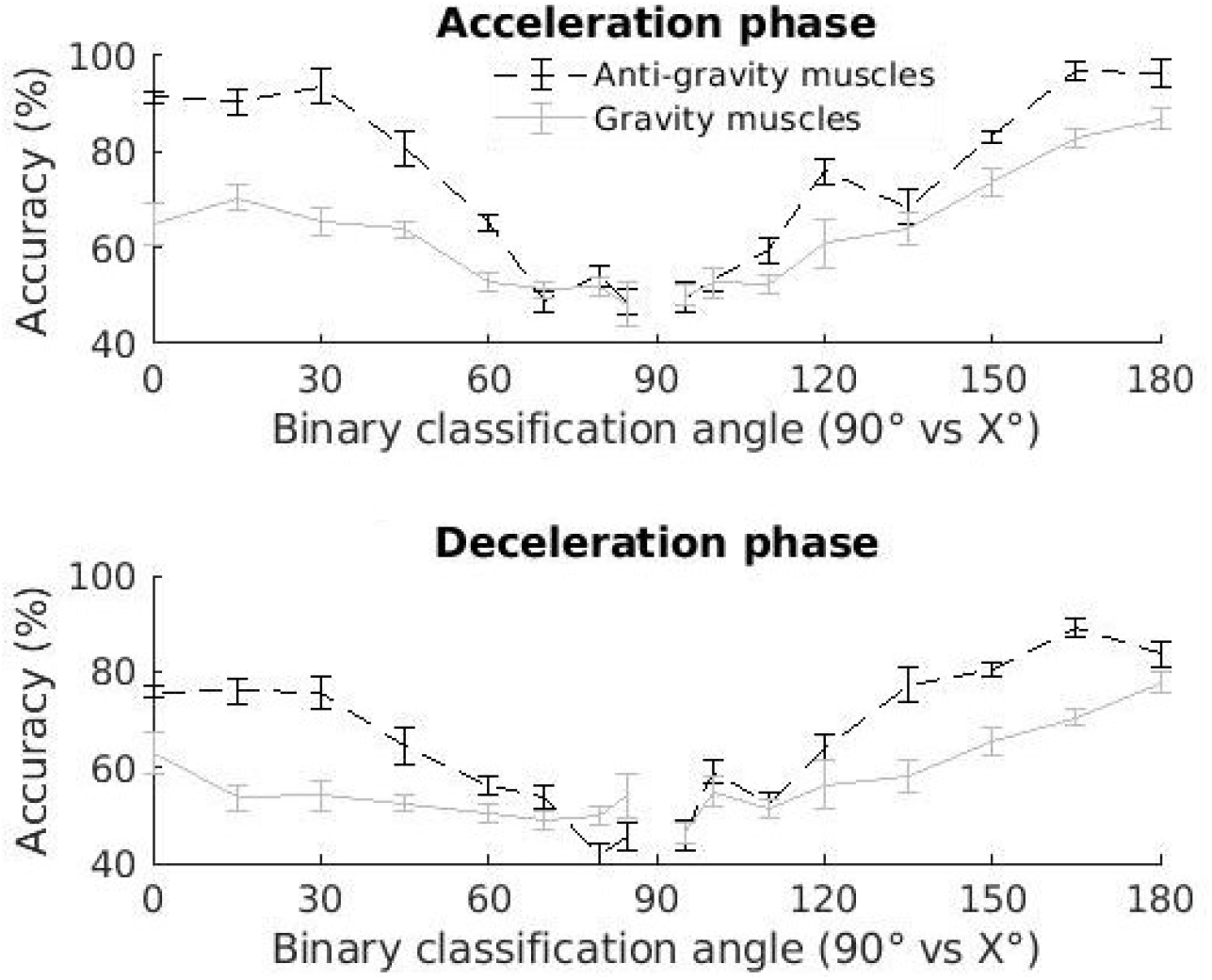
Classification accuracy using only the EMGs from either gravity or antigravity muscles during the acceleration phase of the movement or the deceleration phase, using SVM for classification.

**Figure S6.**
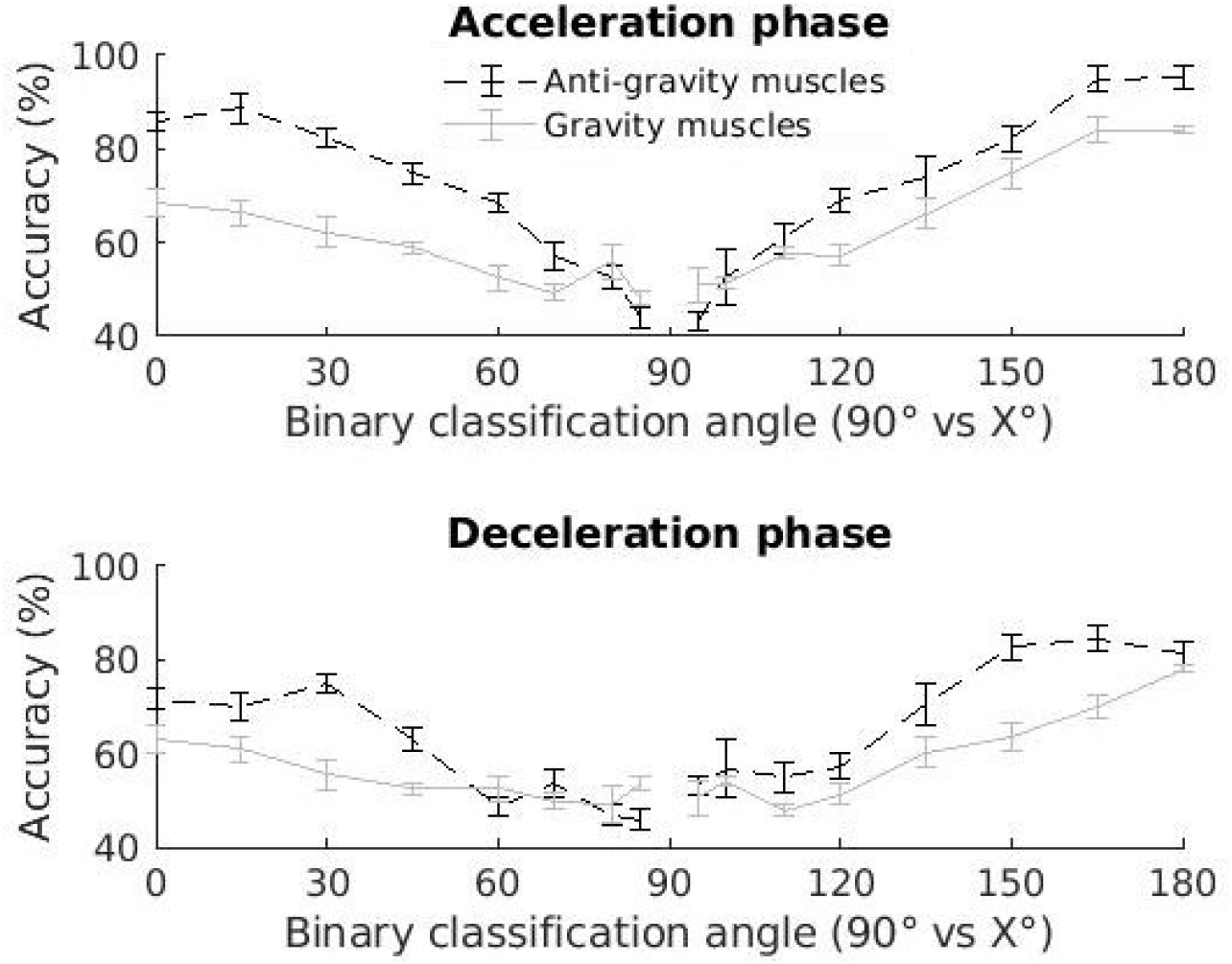
Classification accuracy using only the EMGs from either gravity or antigravity muscles during the acceleration phase of the movement or the deceleration phase, using SVM for classification. Contrary to Figure S5, the biceps brachii is not included in the antigravity muscles here.

**Figure S7.**
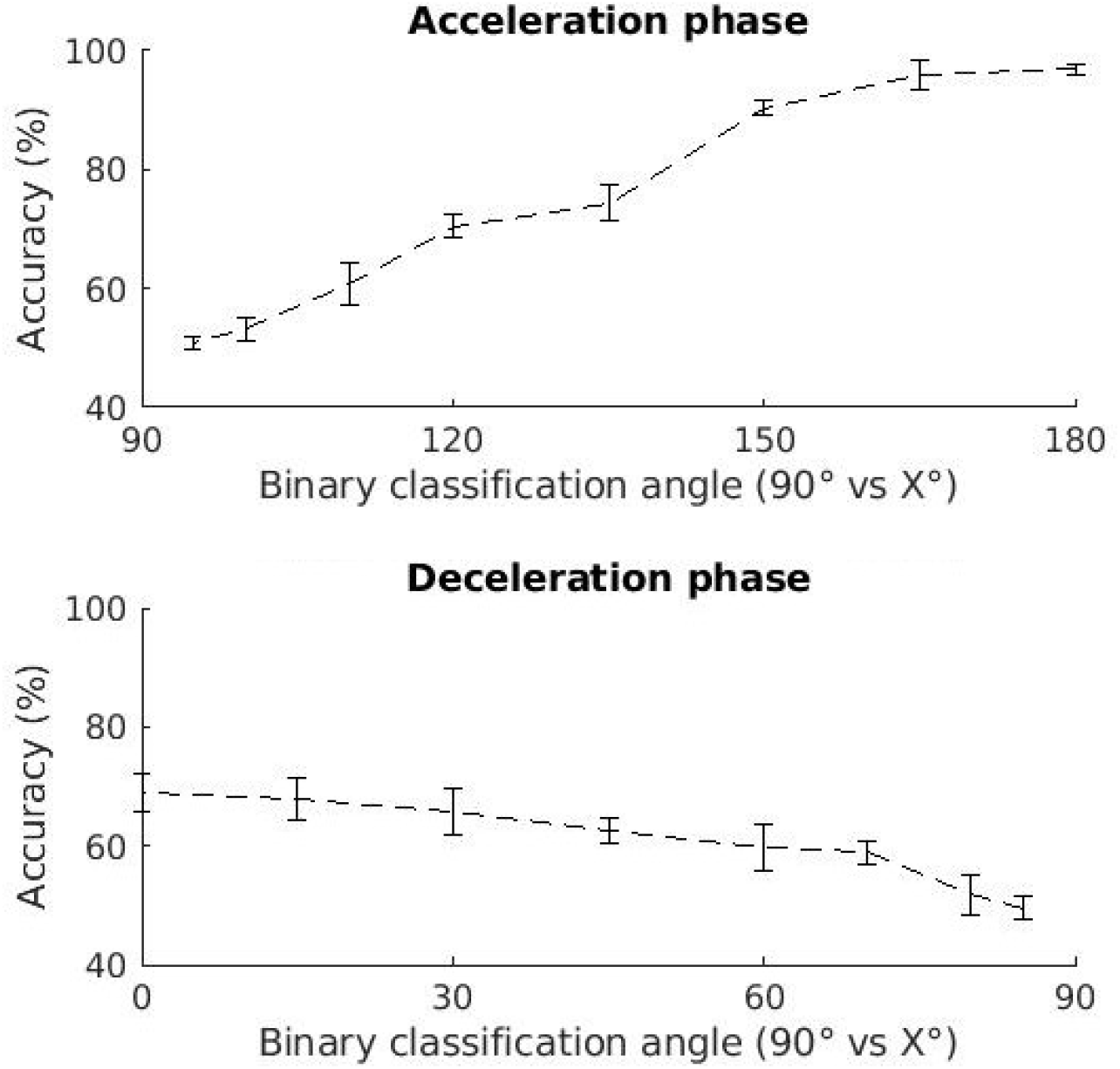
Classification accuracy using only the negative part of the EMGs from antigravity muscles during the acceleration phase of the movement or the deceleration phase, using SVM for classification.

**Figure S8.**
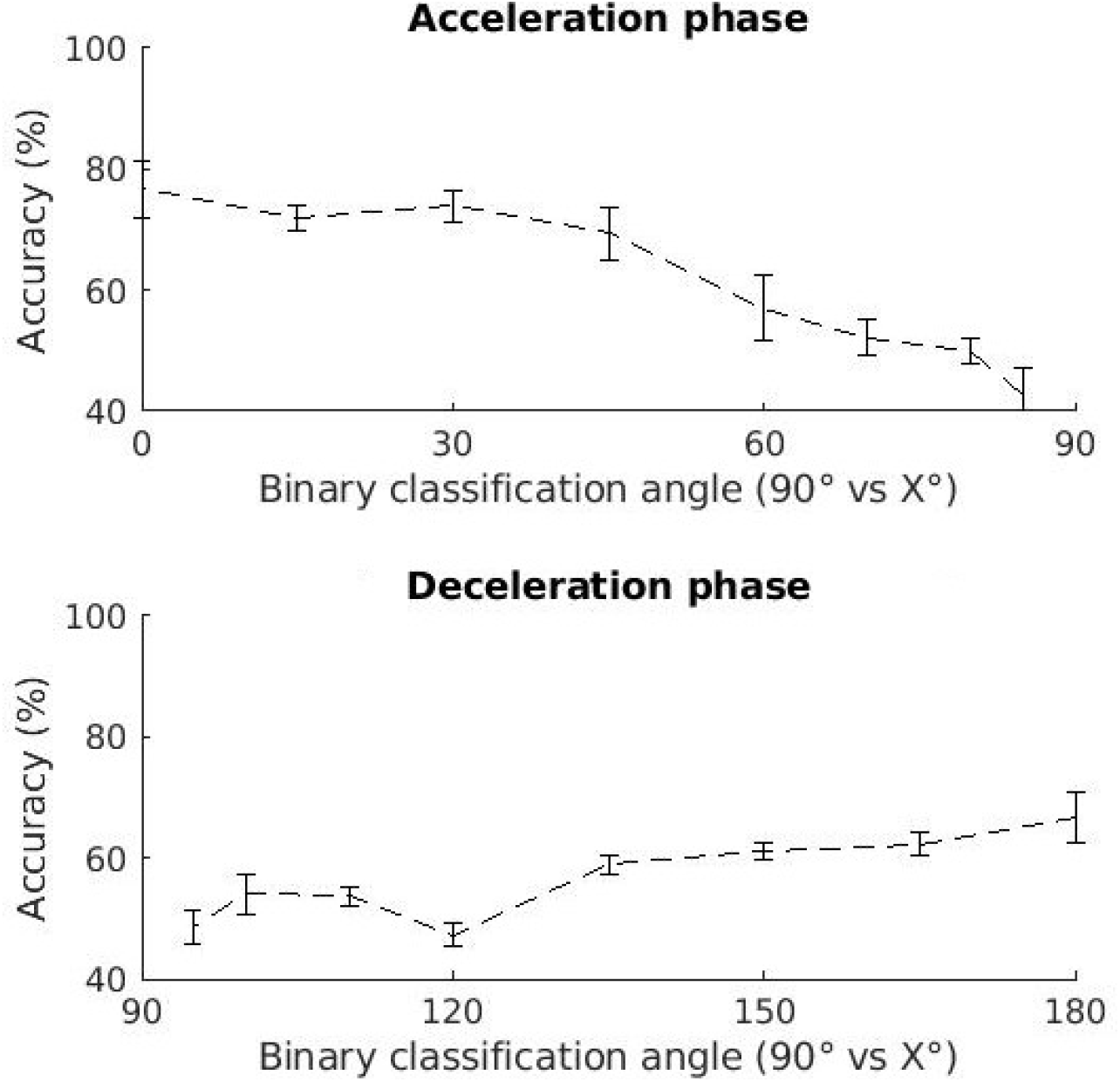
Classification accuracy using only the positive part of the EMGs from antigravity muscles, using LDA for classification.

**Figure S9.**
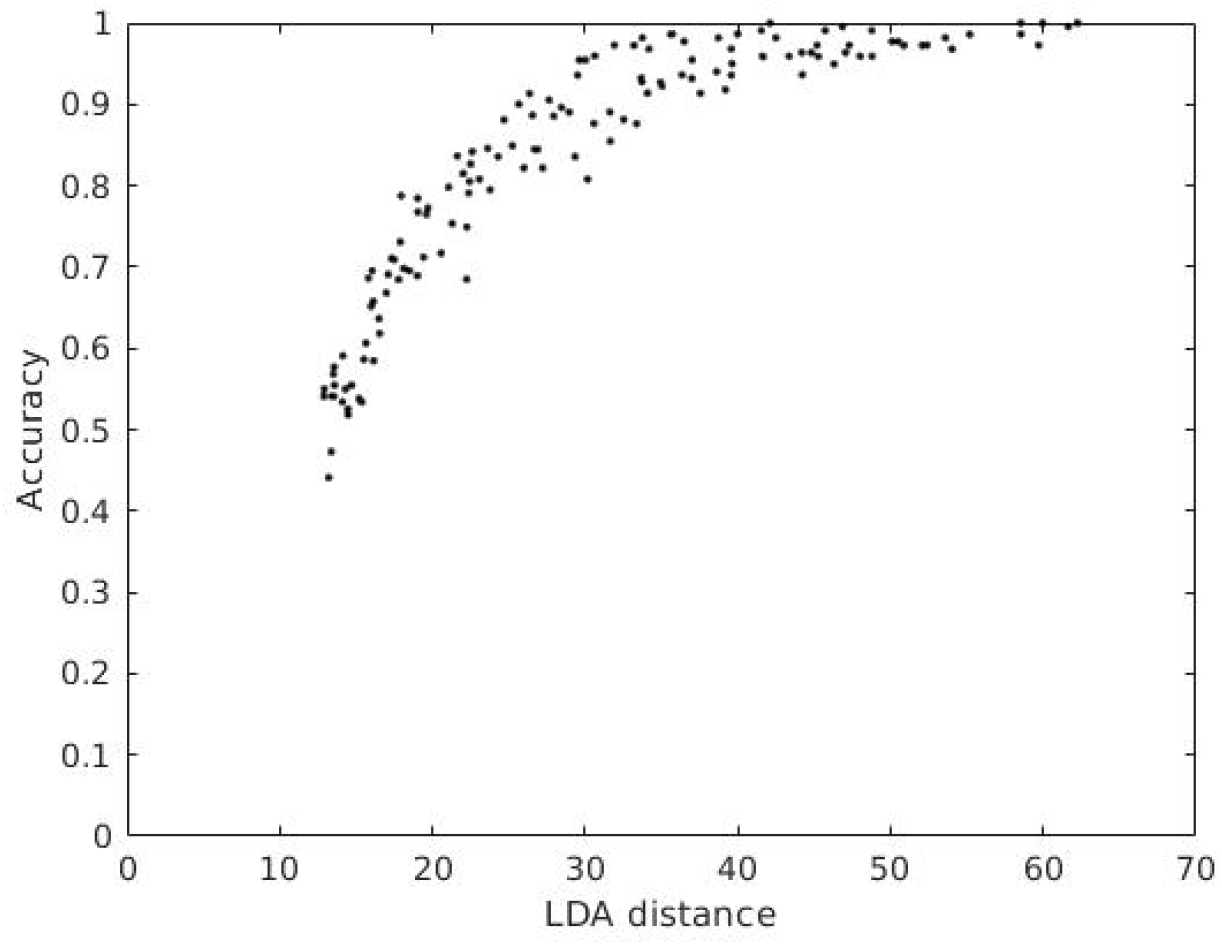
Accuracy as a function of the LDA_distance_.

